# Extensive genetic diversity in *Plasmodium vivax* from Sudan and its genetic relationships with other geographical isolates

**DOI:** 10.1101/2020.06.01.127423

**Authors:** Musab M Ali. Albsheer, Eyoab Iyasu Gebremeskel, Daniel Kepple, Eugenia Lo, Virginie Rougeron, Muntaser E. Ibrahim, Muzamil M. Abdel Hamid

## Abstract

*Plasmodium vivax* malaria is a neglected tropical disease in Africa due to low occurrence rates and lack of accurate diagnosis. Recently, there has been a dramatic increase in *P. vivax* cases in East Africa and reportedly spreading to western countries. This study investigated the geographical origin and genetic diversity of *P. vivax* in Sudan by 14 microsatellite markers. A total of 113 clinical *P. vivax* samples were collected from two districts, New Halfa and Khartoum in Sudan. In addition, data from 841 geographical samples retrieved from the database for global genetic analysis were included in the analysis to further the genetic relationships among the *P. vivax* isolates at regional and worldwide scales. On a regional scale, we observed 91 unique and 8 shared haplotypes amongst the Sudan samples. Such a high genetic diversity compared to other geographical isolates lends support to hypothesis that *P. vivax* was originated from Africa. On a global scale, as already demonstrated, we observed distinct genetic clustering of *P. vivax* isolates from Africa, South America, and Asia (including Papua New Guinea and Solomon Island) with limited admixture in all three clusters. The principal component analysis and phylogenetic tree showed similar clustering patterns and highlighted the contribution of the African isolates to the genetic variation observed globally. The East African *P. vivax* showed similarity with some of the Asian isolates suggesting potential recent introductions. Our results show extensive genetic diversity co-occurring with significant multi-locus linkage disequilibrium, demonstrating the effectiveness of using microsatellite markers to implement effective control measures.

## Background

*Plasmodium vivax* malaria is a neglected tropical disease, and is regarded as a benign and self-limiting infection despite the increasing evidence that the overall burden, economic impact, and the severity of disease caused by *P. vivax* is on the rise [1,2]. Malaria caused by *P. vivax* is posing a significant re-emerging disease threat in many African, South American, and Asian countries caused by both resistance to available therapies [3–7] and the parasite’s unique ability to form hypnozoites, a mechanism that enables *P. vivax* to hibernate and relapse in the liver [8–10]. Moreover, the current knowledge on genetic diversity and population structure of *P. vivax* lags behind that of *P. falciparum* due to lack of appropriate genetic markers for the *P. vivax* genome [11] and its inability to be cultured for long-term in the laboratory except in non-human primates [12, 13] further complicating control efforts. The complete sequencing of the *P. vivax* genome in 2008 provided a valuable resource and added a stimulus to the much-needed study of this neglected parasite [11] and subsequent studies [13] revealed greater genetic diversity of *P. vivax* in comparison to *P. falciparum*.

Differences in *Anopheles* mosquito dynamics provide *P. vivax* with a broader temperature tolerance and ability to be transmitted in climates not tolerated by *P. falciparum* [12, 15–18]. Studies conducted from different geographical regions showed distinctive phenotypic features of the parasite populations, which might be attributed to the presence of barriers to gene flow or strong regional selections [18]. Therefore, analyzing the genetic make-up of the isolates not only would be useful in identifying the geographic origins of these parasites [20–22], but also facilitate meaningful parasite-specific control and preventive measures to be implemented.

To understand the geographical origin and genetic diversity of isolates, microsatellite markers (MS) provide an effective means to determine population structure and for high-throughput genotyping due to their higher polymorphic nature [23–28]. The polymorphic nature of MS loci also permits increased detection of multiclonal infections [28], which can be useful when describing the history of endemicity and the stability of transmission of malaria parasites within a specific global region [29], and can describe infection dynamics *P. vivax* across time [31, 32]. A recent study conducted in Sudan [2] showed a rise in *P. vivax* prevalence with concerning proportions and spreading to new areas including east and central African countries [16], posing new challenges to malaria treatment and control in Sudan. The overall prevalence of *P. vivax* among the malaria cases was 26.6%; of the suspected cases, the proportion of positive results for *P. vivax* was 13.3%, showing significant variations between the states and residence status [2], reflecting the stability of the transmission.

In line with the growing number of *P. vivax* cases, this study analyzed the genetic diversity and population structure of *P. vivax* isolates from Sudan using 14 highly polymorphic tri and tetranucleotide microsatellite markers [30]. An additional 841 global samples were included in the analysis to further understand the genetic origin and diversity of the isolates at a regional and global level. Microsatellite analyses resulted into high number of unique haplotypes and clustering of the isolates into geographical origins.

Genetic analysis of *P. vivax* using microsatellites is robust to address questions on the geographical origin and spread of the parasites and contribute to the current malaria control and elimination effort in Sudan.

## Materials and Methods

### Sample collection and ethics statement

*P. vivax* microsatellite typing was conducted on 113 field isolates collected from two districts in Sudan: 21 from Halfa (2013) and 92 from Khartoum (2013 and 2015). A field isolate is defined in this study as a sample of parasites derived from a single-infected patient collected on a single occasion (19). All the patients were febrile with 37.5°C or above and were diagnosed as having *P. vivax* malaria by microscopic examination of Giemsa-stained thick and thin blood smears. Finger-prick blood samples, for molecular analysis, were collected on Whatman 3 mm filter paper (Whatman International, Ltd., Maidstone, England), dried, and kept in a dark place. Filter paper samples were coded to keep the anonymity of the patients and kept at Parasitology and Medical Entomology laboratory at the Institute of Endemic Diseases, the University of Khartoum for further analysis. Samples were obtained and analyzed after approval from the Ethics Committee of the Institute of Endemic Diseases, University of Khartoum (Reference number: 9/2016).

### Microsatellite typing of *P. vivax*

DNA from blood samples was extracted using DNeasy Blood and Tissue kit (Qiagen, France) according to manufacturer’s recommendations, and eluted in 100 µl of elution buffer per 200 µl of whole blood or per filter plot. For each DNA sample, whole genome amplification was carried out using an Illustra-GenomiPhi™ V_2_ DNA Amplification Kit (GE Healthcare, Uppsala, Sweden) following the manufacturer’s protocols. The product (DNA + Sample buffer) was denatured by heating at 95°C for 3 min and then kept on ice. 9µL of Reaction Buffer (Illustra™ GenomiPhi™ V2 DNA Amplification Kit - GE Healthcare, Uppsala, Sweden) and 1µL of enzyme Phi 29 was added and incubated at 30°C for 2h for genome amplification. The amplification was stopped by placing the samples at 65°C for 10 min, before being stored at -20°C. The product DNA was used as a template for the PCR-based amplification of 14 polymorphic microsatellite markers distributed among 10 of the 14 chromosomes of *P. vivax*. The PCR protocol was adapted from Karunawera et al. (30) as follows: a 20 µl reaction mix was made of 0.3 µl of each primer (10 µM), with the forward being labelled with a fluorochrome, 0.1 µl template DNA, 0.5 µl dNTP mix (10 pmol/µl), 0.7 µl of MgCl_2_ (50mM), 2.5 of 10X buffer, 0.2 µl Taq polymerase (5 UI/µl, Invitrogen). The fourteen highly polymorphic microsatellite markers used in the study for typing consist of tri-nucleotide tandem repeat motifs (MS1, MS6, MS10, MS8, MS9, MS20, MS16, MS2, MS5, MS12, MS15, MS3, MS4, and MS7) and tetra-nucleotide repeat array (MS2) (30).

Amplifications were carried out in a thermal cycler: 40 cycles of denaturation at 94°C for the 30s, the annealing temperature of each locus for 1 min at 58°C and extension at 72°C for 30s, followed by a final extension at 72°C for 7 min. The reaction products were visualized on a 1.5% agarose gel stained with EZ-vision. Fluorescence-labeled PCR products were sized on Applied Biosystems ABI3500XL (GenSeq platform, Montpellier), with a Genescan 500 LIZ internal size standard.

### Data analyses

In addition to the 113 *P. vivax* samples from Sudan, 841 global samples were retrieved from the database for global genetic analysis (http://plasmodb.org Accessed November 12, 2008) to analyze the ancestry of the isolates and to have a regional and global perspective of the parasites studied following the protocols followed in [31]. To maintain consistency, only 11 loci of the 17 populations were considered for analysis.

As blood-stage malaria parasites are haploid, the presence of one or more additional alleles at a particular locus was interpreted as a co-infection with two or more genetically distinct clones (multiple clone infections) in the same isolate [20, 34, 35]. Therefore in co-infection cases, the single or predominant allele at each locus was considered for computing allele frequencies and to define haplotypes [23–28, 36]. The genetic diversity of the parasite populations from Sudan was determined by calculating the heterozygosity (*H*_*E*_) at each locus in each separate population. Virtual heterozygosity is the average probability that a pair of alleles randomly obtained from the population is different and was defined as *H*_*E*_= [*n* /(*n* − 1)][1 − Σ *p* 2 *i*], with *n* is the number of isolates analyzed and *pi* the frequency of the *ith* allele in the population. Virtual heterozygosity ranges between 0 and 1, with values close to 1 reflecting high genetic diversity levels in a population.

Data were also analyzed using LIAN software version 3.5 [35] to calculate a standardized index of association (I^S^_A_) to test for evidence of overall multilocus linkage disequilibrium. This test compares the variance (V_D_) of the number of alleles shared between all pairs of haplotypes observed in the population with the variance expected under random association of alleles (V_E_) and is defined as I^S^_A_ = (V_D_/V_E_ − 1)(r − 1), with r being the number of loci analysed [36]. V_E_ is derived from 10,000 simulated data sets in which alleles were randomly reshuffled among haplotypes. Significant linkage disequilibrium is detected if V_D_ is > 95% of the values derived from the reshuffled data sets [37]. Moreover, GenAlEx program version 6.5 [36,37] was used to estimate allele richness and heterozygosity.

The Bayesian clustering model to assign isolates to *K* populations according to allele frequencies at each locus was applied using STRUCTURE 2.3.1 software [40] to test whether microsatellite haplotypes clustered according to the geographic origins of the isolates. The program was run 5 times at each of 4 different *K* values (*K* 2–6) with a burn-in period of 50,000 iterations followed by 1×10^5^ iterations. The admixture model was used in all analyses. Isolates with predominant ancestry (> 70%) were considered as members of that particular population [19]. PAST software [41] was used to determine the genetic variation among the populations. Phylogenetic tree of the 17 populations from this study and global samples was constructed using GenoDive v. 3.0 [42]. Moreover AMOVA analysis implemented in GENAlex program [38] was conducted to determine the distribution of genetic variation within and among populations included in the study.

## RESULTS

### Microsatellite variation in *P. vivax* isolates from Sudan

All parasite isolates from New Halfa and Khartoum were highly polymorphic and showed 68 unique and 12 shared haplotypes (Supplementary Table 2), with the number of alleles per locus (Na) ranging from 5 to 13 in New Halfa and 6 to 21 in Khartoum (Table 1). The number of alleles observed in isolates from Khartoum is more than the number of alleles observed in isolates from New Halfa (Table 1). The most diverse alleles observed in isolates from New Halfa (0.895) and Khartoum (0.870) is locus MS15 (Table 1). High number of private alleles shared among both populations (Supplementary Figure 1). The virtual heterozygosity (*H*_*E*_) values (Table 2) for the isolates from both New Halfa and Khartoum pooled together ranged from 0.70275 to 0.8931 (mean 0.80293 ± 0.06557). Significant multilocus linkage disequilibrium (Table 3) was found in the isolates (n = 113) from Sudan (*I*_*S*_^*A*^= 0.1486, *P* < 0.001). The association was high (Table 3) when isolates from Khartoum (*I*_*S*_^*A*^= 0.1967, *P* < 0.01) were analysed separately.

**Figure 1.**
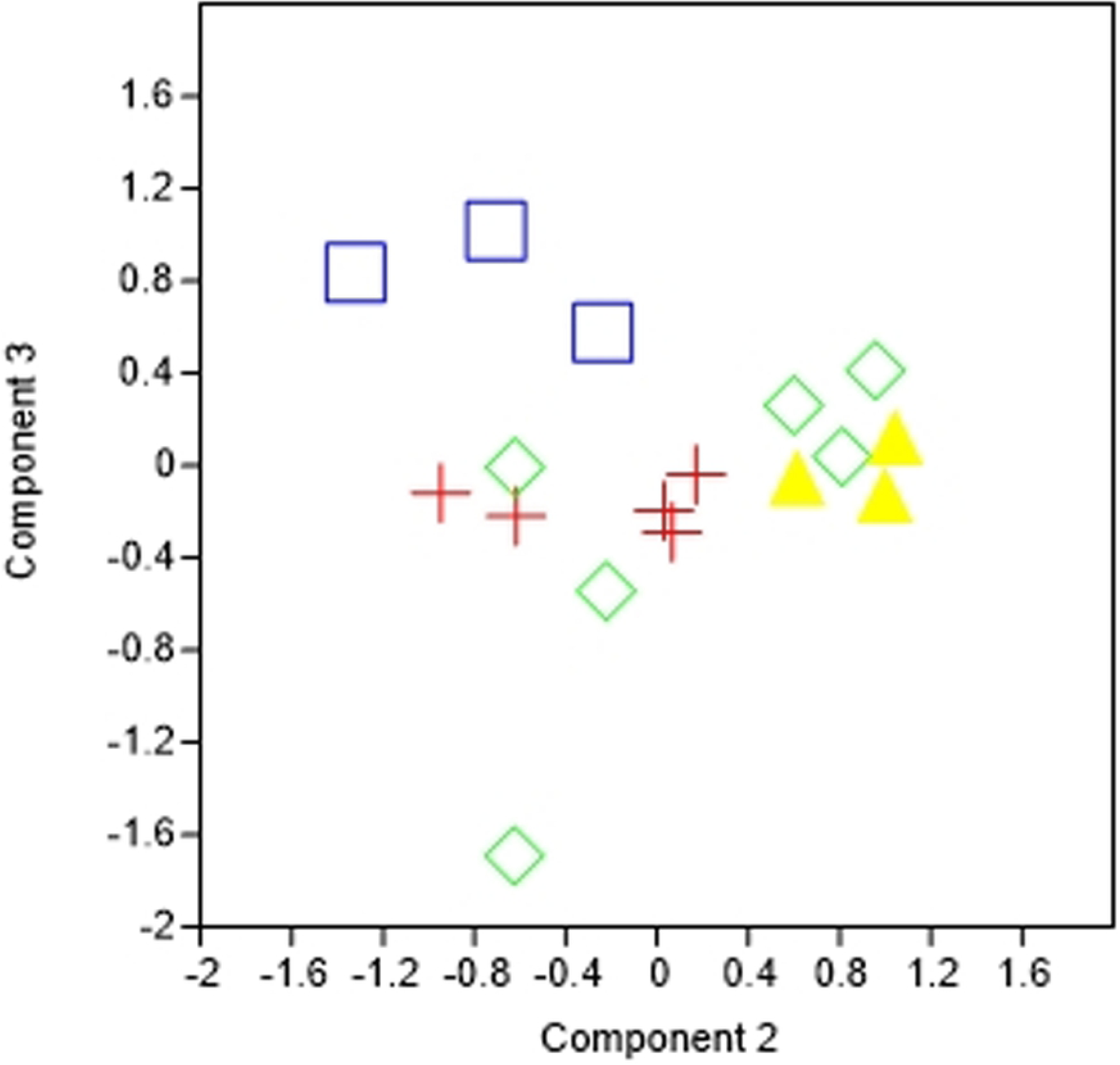
Population structure of 17 populations analysed at the K=2, 3, 4, 5 and 6. Cluster 1 (Red) includes *P. vivax* isolates from Peru, Brazil, and Central America. Cluster 2 (Green) includes *P. vivax* isolates from New Halfa, Khartoum, other parts of Sudan and Africa, Madagascar and isolates from Peru. Cluster 3 (Blue) includes *P. vivax* isolates from Cambodia, Vietnam, Indonesia, Papua New Guinea low land, Papua New Guinea high land and Solomon Island.

### Geographic structure of *P. vivax*

The clustering patterns obtained with *K =3* (Figure 1) showed clear structure of *P. vivax* isolates from Africa, South America, and Asia (including Papua New Guinea and Solomon Island) with much admixture in all three clusters. Cluster 1 (Red) included *P. vivax* isolates from Peru (90.8%), Brazil (95.4%), Central America (50.3%). Cluster 2 (Green) included *P. vivax* isolates from New Halfa (84.5%), Khartoum (81.5%), other parts of Sudan (89.9%), and of other African countries (77.3%), including Madagascar (77.4%). Cluster 3 (Blue) included *P. vivax* isolates from Cambodia (54.4%), Vietnam (76.4%), Indonesia (51.2%), Papua New Guinea low land (92.5%), Papua New Guinea high land (89.4%), and Solomon Island (79.9%). Isolates from the rest three Asian countries Armenia and India were assigned to cluster 2 with 87.5% and 74.4% proportion of the membership, respectively, whereas isolates from Uzbekistan were predominantly present in cluster 1 with 77.5% proportion of the membership. The proportion of the membership of each pre-defined population in each of the three clusters was higher than 75% except for Central America and Indonesia. When the number of clusters was set to 4 or 5 (Figure 1), the separation between countries was limited except for the isolates from Khartoum that formed a separate cluster from isolates from New Halfa and other isolates from Africa. The New Halfa and other isolates from Africa clustered with those from Peru and Uzbekistan

Principal component analysis (PCA) (Figure 2) generated using GenoDive v. 3.0 (40) showed clear clustering of all the populations analyzed into their geographical location to corroborate population structure results observed at K=3 level (Figure 1B). In general, PCA, NJ tree and clustering analyses showed similar structuring patterns. The phylogenetic tree (Figure 3) constructed using PAST further highlights the importance of isolates from Africa and Sudan to the genetic variation observed globally. AMOVA (Table 4) analysis of the 17 populations showed that most of the variation lies within the population (89%) and 11% variation observed among the populations.

**Figure 2.**
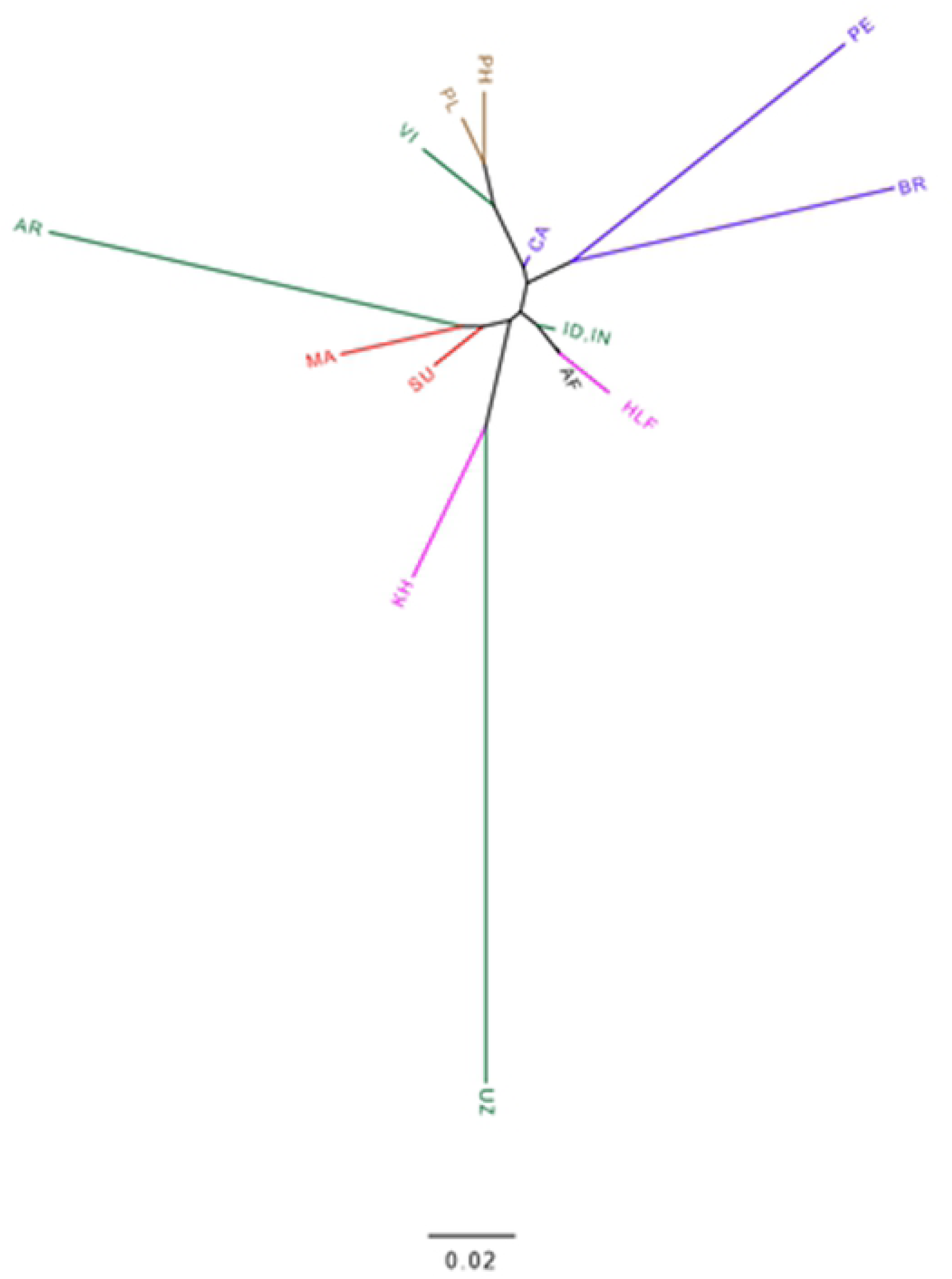
Principal component analysis on Sudanese and global *P. vivax* populations constructed using PAST software based on the genetic distance data generated from GenAlex software. Red includes isolates from New Halfa, Khartoum, Madagascar and isolates from another part of Sudan and Africa; Green includes isolates from Asia (Armenia, India, Cambodia, Vietnam, Indonesia, and Uzbek); Yellow includes isolates from Papua New Guinea and the Solomon Islands; and Blue includes isolates from Brazil, Central America, and Peru.

**Figure 3.**
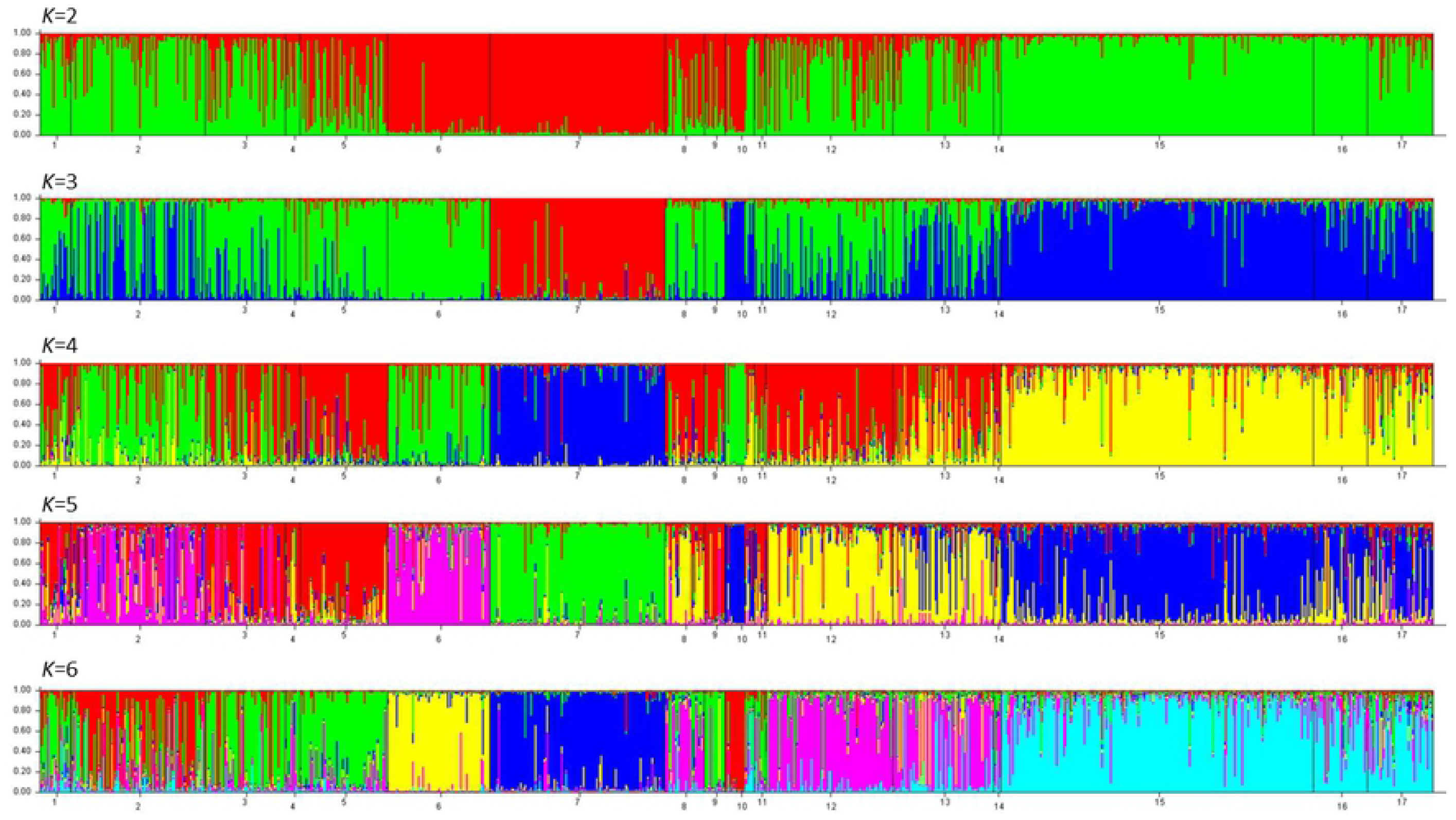
Phylogenetic tree generated for Sudanese and global *P. vivax* populations constructed using GenoDive v. 3.0 [43] based on the genetic distance data generated from GenAlex software. AR: Armenia, BR: Brazil, CA: Central Americas, HLF: New Halfa, Id and In: Indonesia/India, KH: Khartoum, Ma: Madagascar, Pe: Peru, PH: Papua New Guinea highlands, PL: Papua New Guinea Lowlands, Su: Other Sites in Sudan, Uz: Uzmbek, and Vi: Vietnam.

## Discussion

*P. vivax* cases in Sudan rose five times in six years [43]. The rise and re-emergence of *P. vivax* infectious could be related to the parasite’s ability to develop alternative mechanisms to invade human erythrocytes other than the Duffy antigen [44], which also is associated with severe cases once more highlighting of undertaking more studies of the parasite which once was thought as benign and neglected. Continentally, the Duffy negative phenotype ranges from 60% in the eastern countries to 98% in the western countries [45] suggesting a prolonged period of host-parasite co-evolution [46]. Genomic analyses have identified that the evolution of Duffy negative infections occurred independently in a mixed Duffy population [45]. Prior studies have also shown cases of *P. vivax* transmitted from eastern African refugees to natives in Germany and Switzerland [51, 52].

Genetic markers like microsatellites (MS) are selectively neutral and still powerful tools for investigating the population structure parasites because of their polymorphic nature [50]. MS is a popular alternative to polymorphic antigenic genes in studying malaria parasites due to their purported neutral ubiquity throughout genomes and utility for describing the evolutionary history of global populations [51]. Furthermore, the relatively unconstrained polymorphic nature of MS loci permits increased detection of multi-clonal infections [28] which can be useful when describing the history of endemicity and the stability of transmission within a specific global region [29]. In line with this, we tried to characterize the genetic diversity of *P. vivax* populations from blood samples collected in New Halfa and Khartoum in Sudan, by employing MS genotyping and analysis. Overall, in the two populations studied high microsatellite polymorphisms, characterized by high heterozygosity [50] was recorded. This high heterozygosity observed is extended to all individual loci analyzed with mean H_E_ of 0.80293 (SD=0.06557). The observed high level of genetic diversity in low to meso-endemic setting countries like Sudan is not uncommon as it corroborates with other studies [20, 34, 56–59], lending support to the notion that Africa is the origin of *P. vivax* [60, 61] though it is disputed [59].

The use of 11 highly polymorphic microsatellite markers identified population structure of *P. vivax* isolates as Asian and Latin American isolates and the isolates from Africa clustering with SE Asian isolates. With the addition of a third population (*K = 3*), further differentiation between global isolates isolates from Brazil sitting in one cluster and the isolates from the Solomon Islands and Papua New Guinea assuming a separate cluster. When the number of clusters was set to 4 or 5, the separation between countries was limited [23] except for the isolates from Khartoum then assuming a separate cluster from isolates from New Halfa and other isolates from Africa, which the latter clustered with isolates from Peru and Uzbekistan highlighting the importance of designing studies at sub-regional or sub-population level in this part of Africa as was seen in human population studies [58]. The genetic association observed between the East African and Asian isolates may imply a recent re-introduction via human migration of *P. vivax* from Asia to East Africa. With all the limitation of using microsatellites in studying population genetics of parasites [52], the population structure generated using these microsatellites show promise in identifying origin [19] and providing useful information regarding the genetic structure of parasites [61, 62].

### Conclusions

Microsatellite typing remains an important tool to study *P. vivax* population structure on a global scale, and differences in diversity reflect transmission intensity and isolation of parasite populations as was reported elsewhere [23]. Microsatellite-based analysis of *P. vivax* parasites from Sudan showed extensive genetic diversity co-occurring with significant multilocus linkage disequilibrium. These parasite populations were seen to cluster according to their geographic locations and ancestry. As inferred elsewhere the predictive power of ancestry with this microsatellite data as a model is promising and could be useful in identifying the origin of *P. vivax* malaria cases and enabling meaningful control and preventive measures to be implemented. Moreover, dissecting the parasite population into subpopulations and sub-regions especially in this part of the world would give a meaningful interpretation in the origin and structuring of *P. vivax* parasites in the future.

## Acknowledgements

We are greatly indebted to the patients for participating in the study by donating blood and the staffs and technicians from the Institute of Endemic Diseases, University of Khartoum for assisting sample collection; the communities and hospitals for their support and willingness to participate in this research. Authors would like also to thank IRD, CNRS-INEE, and the ANR Tremplin EVAD 2017 for generation of the data.

## Tables

**Table 1**. Isolates from New Halfa (N) and Khartoum (N) were analyzed separately using the GENAlex program to determine allelic diversity in the populations at an individual locus. The table shows sample size (N), number of alleles (Na), number of effective alleles (Ne), information index (I), Diversity (h) and unbiased diversity (uh) for every 11 individual loci.

**Table 2**. All isolates from Sudan (N) were analysed using GENAlex program to determine the number of alleles (Na), the number of effective alleles (Ne), information index (I), Diversity (h), unbiased diversity (uh) and virtual heterozygosity (H_E_) for each locus.

**Table 3**. The genetic diversity of the parasite isolates from New Halfa, Khartoum and all samples from Sudan (all samples pooled) were determined by calculating the virtual heterozygosity (H_E_) using LIAN software.

**Table 4**. AMOVA results showing genetic variation within and among populations implemented in GENAlex program. The P value is 0.001.

Supplementary Files

**Supplementary Figure 1**. Estimate of allele richness and private alleles in both populations analysed. Data generated using GenAlEx program.

**Supplementary Table 1**. Haplotypes generated using Arlequin program [63] with 68 unique haplotypes and 12 shared haplotypes in all the isolates from this study.

